# Long-term mitigation of the foreign-body response with dexamethasone-eluting cochlear implants in mice

**DOI:** 10.64898/2026.08.26.747195

**Authors:** Aditya Alluri, Bryce Hunger, Fahad Hossain, Shakila Mahmuda Fatima, Muhammad Taifur Rahman, Brian J. Mostaert, Robert Gay, Ya Lang Enke, Marlan R. Hansen, Alexander D. Claussen

**Author notes:** **Corresponding Author:** Alexander D. Claussen, Department of Otolaryngology-Head and Neck Surgery, University of Iowa, Iowa City, IA-52242, USA. Authors contributed equally to work as co-first authorship (Aditya Alluri, Bryce Hunger).

## Abstract

The inflammatory foreign body response that follows cochlear implantation produces intracochlear fibrosis, neo-ossification, and elevated electrode impedances that can compromise implant performance. Dexamethasone-eluting cochlear implants reduce this response, but the durability of their anti-inflammatory effect over long implantation intervals has not been established. Using a murine model of chronic cochlear implantation in CX3CR1+/eGFP Thy1+/eYFP dual-reporter mice, we compared dexamethasone-eluting and standard mouse cochlear implants at 224 and 336 days post-implantation. Density of CX3CR1+ macrophages, MHCII+CX3CR1+ antigen-presenting macrophages, α-SMA+ fibrosis, and neo-ossification were quantified in the scala tympani, Rosenthal canal, and lateral wall of the basal turn. Standard implants produced persistent macrophage and antigen-presenting macrophage infiltration, accompanied by an α-SMA+ fibrotic response and neo-ossification. Dexamethasone-eluting implants suppressed macrophage infiltration in all three regions out to 336 days and reduced fibrosis at 224 days. In the subset of cochleae with electrode array translocation, dexamethasone-eluting implants attenuated macrophage infiltration and confined the fibrotic and osseous response to the site of translocation, whereas standard implants produced a widespread response. A reduction in immune cell density was also observed in the contralateral, unimplanted cochleae of animals implanted with dexamethasone-eluting implants, suggesting a wider component to the drug’s effect. Dexamethasone-eluting cochlear implants therefore provide sustained, long-term suppression of the cochlear foreign body response in mice, supporting their continued translation toward clinical application. This effect was associated with continued low-level dexamethasone elution out to 336 days post-implantation; further work is needed to assess the durability of this effect at the conclusion of drug elution.

**Highlights:**

- Dexamethasone-eluting implants suppress cochlear foreign body response out to 336 days
- Low-level dexamethasone elution persisted through 336 days post-implantation
- Dexamethasone-eluting implants reduce inflammatory response to array translocation

## 1. Introduction

Hearing loss is a common sensory deficit thought to affect around 30% of the global adult population (Tao et al., 2025). According to the 2019 Global Burden of Disease study, hearing loss was ranked as the third leading cause of years lived with disability and was the leading cause of years lived with disability among sensory losses. The prevalence of hearing loss continues to grow as the population ages, and by 2050, a projected 2.45 billion people will suffer from hearing loss (Haile et al., 2021). In addition to directly affecting the perception of sound, hearing loss negatively impacts children’s language development, is a modifiable risk factor for dementia, and often contributes to social isolation and conditions such as anxiety and depression (Shukla et al., 2020; Mener et al., 2013; Contrera et al., 2017; Lieu et al., 2020; Loughrey, 2025).

Cochlear implants (CIs) are commonly used neuroprosthetic devices that help restore hearing perception in individuals with sensorineural hearing loss. As of July 2022, more than one million CIs have been implanted worldwide (Zeng, 2022). These devices function by bypassing cochlear hair cells and directly stimulating spiral ganglion neurons (SGNs) with an electrode to provide the sensation of sound. Many different technological advancements, such as those in CI design, surgical technique, and processor programming, have occurred (Mitchell-Innes et al., 2018; Roche and Hansen, 2015), allowing for preservation of residual acoustic hearing and improvement in key outcomes such as speech understanding in noise, sound localization, and music appreciation (Shukla et al., 2020; Mener et al., 2013; Contrera et al., 2017).

Although CIs are generally considered to be safe and biocompatible, cochlear implantation triggers an inflammatory response followed by fibrotic tissue formation around the electrode array. This response results in a dense fibrotic sheath directly surrounding the electrode array, as well as looser fibrotic tissue, granuloma formation, and neo-ossification in nearby areas of the cochlea (Foggia et al., 2019; Marsh et al., 1992; Nadol et al., 2001; Schindler and Bjorkroth, 1979; Benatti et al., 2013; Seyyedi and Nadol, 2014; Quesnel et al., 2016; Linthicum et al., 1991; Li et al., 2007; Fayad et al., 2009; Ishai et al., 2017; Rahman et al., 2022). The presence of this response is nearly ubiquitous in the cochlea after cochlear implantation (Linthicum et al., 1991; Li et al., 2007). The tissue response is most robust in the basal turn of the scala tympani (ST) in the region directly adjacent to the implant, but has also been seen distal to the implant, in other scalae, and in the lateral wall of the cochlea (Foggia et al., 2019; Quesnel et al., 2016; Linthicum et al., 1991; Li et al., 2007; Fayad et al., 2009; Ishai et al., 2017; Rahman et al., 2022). This tissue response has been shown to be associated with numerous clinically important negative outcomes, such as poorer word recognition scores (Kamakura and Nadol, 2016) and loss of residual acoustic hearing (Quesnel et al., 2016; Scheperle et al., 2017; Tejani et al., 2022).

The post-implantation tissue response comprises both immediate and delayed components. The immediate tissue response is thought to be a result of traumatic insertion of the CI, whereas the delayed tissue response is thought to be due to a foreign body response (FBR) to the CI materials, as the electrode arrays are considered biocompatible, but are not bioinert (Foggia et al., 2019; Rahman et al., 2022; Rahman et al., 2023; Jensen et al., 2022; Seyyedi and Nadol, 2014). This delayed FBR, along with inner hair cell loss, is a contributor to the phenomenon of delayed hearing loss after implantation, which has been shown to affect 30-40% of patients with initially preserved acoustic hearing (Gantz et al., 2018; O’Malley et al., 2024; Wu et al., 2026; Jensen et al., 2022).

Macrophages infiltrate the cochlea within days of implantation and persist chronically, contributing to a sustained FBR (Gantz et al., 2018; Okayasu et al., 2020; O’Malley et al., 2016; Claussen et al., 2019; Irving et al., 2013; Mistry et al., 2014). In addition, alpha-smooth muscle actin (α-SMA) and type 1 collagen, markers of contractile myofibroblasts and mature fibrosis, have been detected in the fibrotic response over weeks to months (Bas et al., 2015; Hinz et al., 2001; Trojanowska et al., 1998).

Dexamethasone, a synthetic glucocorticoid, exerts anti-inflammatory effects through binding of cytoplasmic glucocorticoid receptors, leading to nuclear translocation and transcriptional suppression of pro-inflammatory cytokines (IL-1β, TNF-α, IL-6), inhibition of macrophage recruitment and activation, and direct suppression of fibroblast proliferation and collagen synthesis (Cain and Cidlowski, 2017). Intravenous use preoperatively or local application in the round window niche perioperatively has been shown to reduce impedance clinically and protect residual acoustic hearing in animal models (Ardıç et al., 2023; Lo et al., 2017; James et al., 2008). Recent advances include dexamethasone-eluting CIs (Dex-CIs), which provide sustained intracochlear dexamethasone delivery and have been shown to reduce electrical impedance in clinical trials (Briggs et al., 2020; Khan et al., 2026a). In animal models, Dex-CIs have been shown to reduce the FBR and impedances, as well as protect hair cells (Ahmadi et al., 2019; Bas et al., 2019; Bas et al., 2016; Liu et al., 2015; Simoni et al., 2020; Van De Water et al., 2010; Manrique-Huarte et al., 2020).

To further investigate the cellular and molecular mechanisms underlying dexamethasone’s effects, murine models of cochlear implantation have been developed. Claussen et al. (2019) previously described the use of a mouse model for cochlear implantation with electrical stimulation. This is notable, as the immune system of mice has been studied extensively, and it enables the use of the many genetic tools available in mice to further investigate the FBR after cochlear implantation. Using this model, further studies found that chronic cochlear implantation leads to robust CX3CR1+/eGFP macrophage infiltration into the cochlea as well as a fibrotic response associated with rising impedances until 56 days (Claussen et al., 2022a; Rahman et al., 2023).

We recently demonstrated that Dex-CIs suppress the cochlear FBR and reduce electrode impedances through 112 days post-implantation in this murine model (Rahman et al., 2025). Dex-CIs were shown to have a dramatic antifibrotic and anti-inflammatory effect and reduced the FBR and electrical impedance to a far greater extent when compared to locally applied dexamethasone in the round window niche. In addition, Dex-CI use was shown to not impact spiral ganglion neuron survival.

The durability of Dex-CI’s suppressive effects during extended implantation periods with continued low-level drug elution remains unexplored. Here, we extend our prior observations to 224 and 336 days post-implantation to test the hypothesis that sustained low-level dexamethasone elution maintains suppression of macrophage infiltration, antigen-presenting cell activation, and intracochlear fibrosis throughout chronic implantation.

## 2. Materials and methods

### 2.1 Sex of mice

Previous studies in both humans and mice have reported no sex-related differences in tissue response to cochlear implantation or in electrode impedance measures (Claussen et al., 2019; Claussen et al., 2022a). Accordingly, animals of both sexes were included in this study, and data were pooled for analysis.

### 2.2 Animals

All the experimental animal protocols were approved by the University of Iowa Institutional Animal Care and Use Committee, consistent with the Guide for the Care and Use of Laboratory Animals from the Institute for Laboratory Animal Research, National Research Council. Subjects included CX3CR1+/eGFP Thy1+/eYFP 8- to 12- week-old mice with a C57BL/6J/B6 background. This mouse model allows for visualization of CX3CR1+/eGFP macrophages and Thy1+/eYFP spiral ganglion neurons.

### 2.3 Cochlear implantation in murine model

Cochlear implantation was performed in the left ear of the mice using Cochlear Limited (Sydney, Australia) standard (mReg-CI) and dexamethasone-eluting (mDex-CI) non-functional HL03 implants. mDex-CIs contained strips of silicone embedded with dexamethasone on the intracochlear portion of the electrode. The cochlear implantation surgical procedure utilized a round window approach of the left ear of the mice under 1-3% isoflurane anesthesia gas, as previously described (Claussen et al., 2019). Mice were followed for either 224 or 336 days after cochlear implantation (**Fig. 1**).

**Fig. 1.**
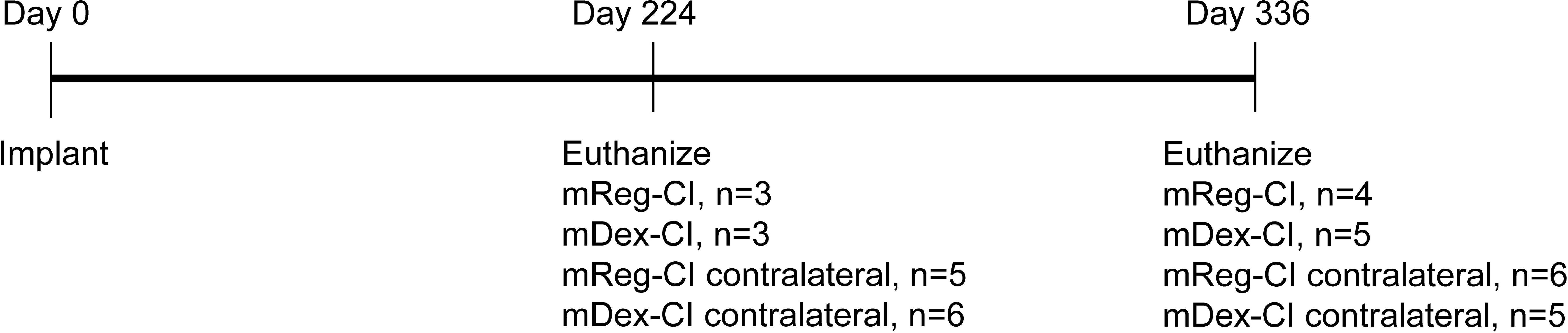
Experimental design depicting the number of subjects in each group and time point with a non-translocated CI that were included in the main analysis.

### 2.4 Dexamethasone content

Cochlear implant devices were explanted post-euthanasia and were analyzed for residual dexamethasone content using a Waters Xevo TQ-S cronos triple quadrupole mass spectrometer with an Acquity UPLC H-Class liquid chromatography system. The liquid chromatography (LC) mobile phases were 25 mM Ammonium Acetate with 0.6% Acetic Acid (v/v) in water (Solvent A) and Acetonitrile (Solvent B). A five-minute isocratic LC separation was performed at 40% Solvent B using a flow rate of 0.4 ml/min. The LC column used was a Waters Acquity BEH C18 (2.1 × 100 mm, 1.7 µm), and it was held at 40°C. The injection volume for each sample was 0.5 µl. The mass spectrometry analysis was performed using positive electrospray ionization (ESI) and multiple reaction monitoring (MRM). The ESI source parameters used were Capillary voltage 1.5kV, Source temperature 150°C, Desolvation Gas (nitrogen) temperature 400°C; Cone Gas flow 50 l/Hr, and Desolvation Gas flow 800 l/Hr. The three MRM transitions [(M+H)+ ion to product ions] were used for quantification: 393.13→373.14, 393.13→355.19, and 393.13→147.13. The cone voltages for each MRM transition were 8V, 26V, and 26V, respectively. The collision energies (eV) used for each transition were 6, 10, and 26, respectively. Waters MassLynx 4.2 software was used for data acquisition and TargetLynx was used for quantitative analysis.

### 2.5 Sample preparation

Prior to perfusion and euthanasia, mice received an intraperitoneal injection of ketamine (80 mg/kg) and xylazine (10 mg/kg). Mice were then transcardially perfused with chilled Phosphate Buffer Solution (PBS) for drainage of blood and then perfused with 4% paraformaldehyde (PFA) as a fixative. Cochleae were extracted and left on a rotator overnight in PFA at 4°C and washed in PBS the following day to remove excess PFA. Cochleae were then decalcified for 5 days by sitting in 0.1 M EDTA (pH 7.5). Samples were then washed in PBS 3 × 10 minutes and cryopreserved using sucrose concentrations 10-30%. After cryopreservation, samples were infused and embedded in optimal cutting temperature medium, mounted to the stage of a sliding block microtome or Leica cryostat and sectioned at a thickness of 30 microns, parallel to the mid-modiolar plane. Sections were mounted on Fisher Superfrost slides and stored at 4°C for 1-7 days before immunolabeling.

### 2.6 Immunohistochemistry

For immunolabeling, slides were first washed 3 × 10 minutes in PBS to wash away excess debris from slides and hydrate the sections. Slides were then washed 3 × 10 minutes in washing buffer (0.1% Triton X-100 and 0.3% Tween 20 in TBS) and then permeabilized and blocked in blocking buffer (1% BSA in washing buffer solution) for 2 hours. Blocked sections were then labeled with primary antibody solutions (Alpha-smooth muscle actin monoclonal antibody, 1A4, eBioscience, catalog# 14-9760-82 (1:1000) and MHC Class II (MHCII) Monoclonal Antibody (M5/114.15.2) eBioscience, catalog# 14-5321-82 (1:200)) in blocking buffer overnight at 4°C with wet KimWipes to keep samples hydrated. The next day, samples were washed 3 × 10 minutes in washing buffer solution and then labeled with secondary antibody solution (Alexa Fluor 568, Invitrogen, catalog# A-11004 (1:400) and Alexa Fluor™ 750, Novus, catalog# NBP2-68490 (1:400)) in blocking buffer for 2 hours at room temperature. Sections were then washed 3 × 10 minutes in washing buffer and then labeled in DAPI (Hoechst 3342 (Sigma) 10 µg/ml) solution in TBS for 30 minutes at room temperature. Slides were then washed 3 × 10 minutes in PBS and subsequently cover-slipped using Epredia Aqua-Mount Slide Mounting Media (catalog# 14-390-5).

### 2.7 Movat’s Pentachrome stain

Slides containing 30-micron-thick cochlear sections were hydrated with distilled water for 5 minutes prior to staining. Sections were then stained with working elastic solution (0.3 ml of Hematoxylin 5%, 0.15 ml of Ferric chloride solution 10%, and 0.15 ml of Lugol’s Iodine Solution) for 20 minutes. The sections were then rinsed in running tap water for 15 minutes followed by 30-40 dips in Ferric chloride differentiating solution.

Next, the slides were washed briefly in tap water followed by 2 changes in distilled water. Sections were then placed in sodium thiosulfate solution (5%), followed by a 2-minute rinse in tap water and two brief changes in distilled water. The slides were placed in 1% acetic acid solution for 2 minutes and then subsequently placed in alcian blue solution for 25 minutes. Next, slides were washed in tap water for 2 minutes followed by 2 brief changes of distilled water. Sections were then stained for 2 minutes in Biebrich scarlet-acid fuchsin solution followed by 2 changes of distilled water. Slides were then washed in 1% acetic acid solution for 20 seconds with continuous agitation followed by a brief rinse in distilled water. Slides were then differentiated in two changes of phosphotungstic acid solution for a total of 22 minutes and then briefly rinsed in distilled water. Slides were then washed in 1% acetic acid solution for 1 minute and then immediately placed in yellow stain solution for 15 minutes. Lastly, slides were rinsed in 3 changes of absolute alcohol and mounted with Permount.

### 2.8 Image acquisition and analysis

Three mid-modiolar cochlear sections per sample were fluorescently imaged on a Leica Stellaris 5 confocal system utilizing the 20 x (0.70 NA) objective, 0.75x digital zoom, and the z-axis spacing set to 1 micron, allowing acquisition of a volumetric z-stack. After image acquisition, images were analyzed using IMARIS (Oxford Instruments, UK).

Volumetric measurements of the scala tympani (ST), Rosenthal canal (RC), and lateral wall (LW) were obtained. Within the ST, CX3CR1+ macrophages and MHCII+CX3CR1+ macrophages were counted, and the volume of α-SMA was quantified. In the LW, CX3CR1+ macrophages and MHCII+CX3CR1+ macrophages were counted. In the RC, CX3CR1+ macrophages, MHCII+CX3CR1+ macrophages, and Thy1+ neurons were counted, and in the middle and apical regions of RC the neurons were counted. Raw counts were then converted to density measurements based on the previously obtained volume of the given regions. Neuron densities of implanted cochleae were compared to their corresponding contralateral ear and assessed as percent neuron loss with the contralateral ear serving as the standard. The fibrotic response was analyzed by volumetric quantification of α-SMA and divided by overall volume of the ST within the section examined to derive a percent of ST occupied by α-SMA.

In an effort to exclude separate inflammatory effects of insertion trauma, subjects with CI scalar translocations were excluded from the main analysis of non-translocated cases and separately analyzed.

Density measurements of immune cells, immune markers, neurons, and fibrotic response in the ST, RC (basal and apical regions), and LW were analyzed in GraphPad Prism. Notably, this main analysis was exclusively performed on subjects without evidence of CI scalar translocation, judged by Movat’s Pentachrome staining. Within GraphPad Prism, two-way ANOVA and Tukey’s multiple comparisons were used to assess differences of interest (α=0.05).

### 2.9 Software

For confocal imaging and image analysis, the most recent version of LAS-X (Leica Microsystems; https://www.leica-microsystems.com/products/microscope-software/p/leica-las-x-ls/) and IMARIS 10.2 (Oxford Instruments; https://imaris.oxinst.com/products/imaris-essentials) were used. Statistical analyses were performed using GraphPad Prism 11 (https://www.graphpad.com/). Adobe Illustrator 25 (Adobe Inc.; https://www.adobe.com/products/illustrator.html) was used for preparation of figures.

## 3. Results

### 3.1 Dexamethasone-eluting CIs continue to elute low levels of dexamethasone through 336 days post-implantation

As shown in **Fig. 2**, analysis of explanted mDex-CIs revealed progressive depletion of dexamethasone content from baseline (day 0) through 224 and 336 days post-implantation, consistent with sustained drug elution. When combined with our previously published data (Rahman et al., 2025), a decrease in residual dexamethasone content in the mDex-CI is observed sequentially at 10, 56, 112, 224, and 336 days post-implantation, corresponding with elution of the dexamethasone content over time. LC– MS/MS analysis detected residual dexamethasone at 336 days post-implantation, as well as a decrease in dexamethasone content from 224 to 336 days post-implantation, suggesting that low-level drug release persists at these time points.

**Fig. 2.**
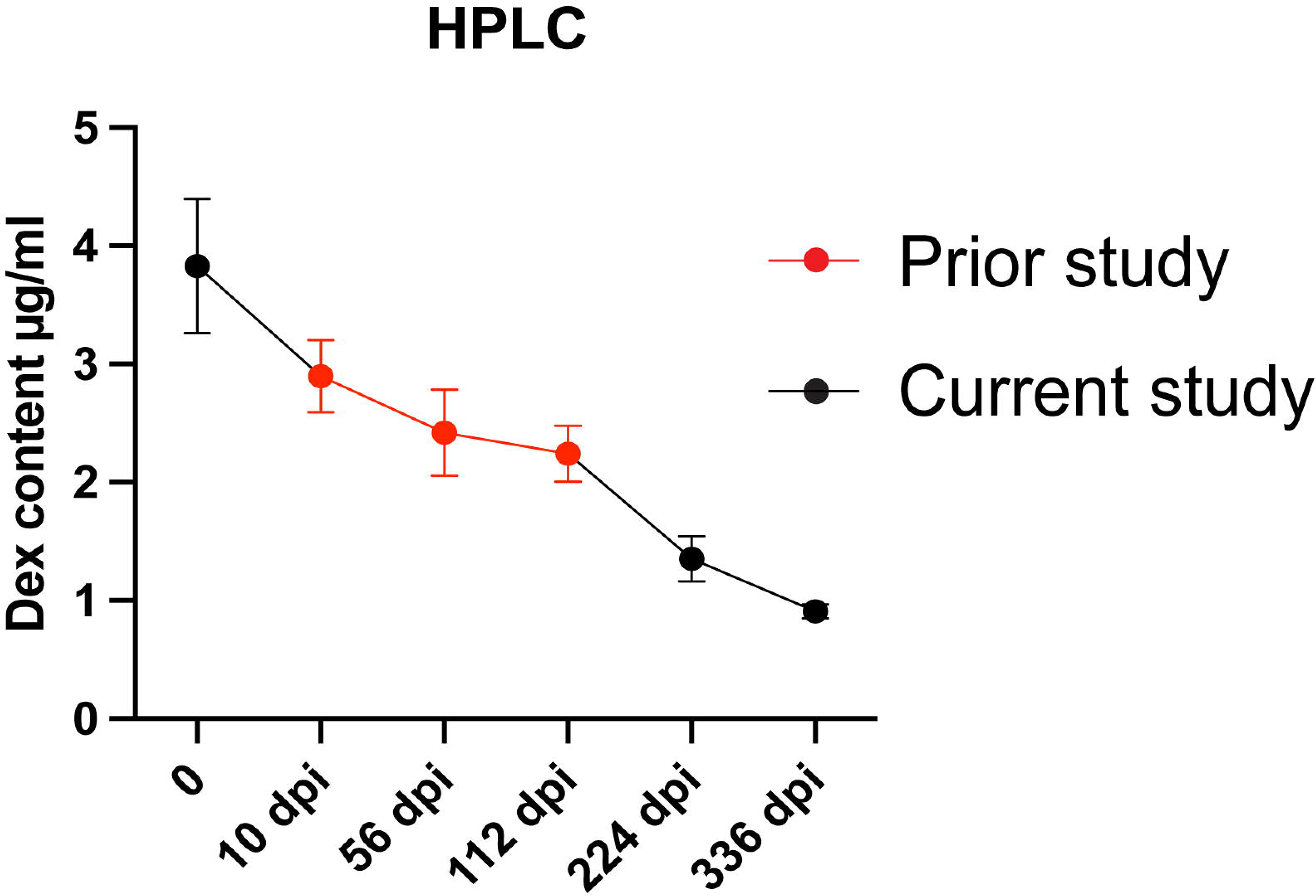
Liquid chromatography–tandem mass spectrometry (LC–MS/MS) analysis of dexamethasone content before implantation (day 0, n=3) and after CI explantation at 224 days (n=6) and 336 days (n=2) post-implantation. Day 0 represents unimplanted mDex-CI. Error bars indicate SEM. Prior study data are from Rahman et al., 2025.

### 3.2 Dexamethasone-eluting CIs reduce FBR in the base of the cochlea

As shown in **Figs. 3**, **4**, mReg-CI implantation results in infiltration of CX3CR1+ macrophages into the cochlea. Compared to mReg-CI, mDex-CI suppresses the infiltration of these macrophages at 224 days post-implantation in the ST (p<0.0001; 12.297 vs 0.190 cells/mm^3^) and LW (p=0.0113; 35.667 vs 9.78 cells/mm^3^) in the basal turn of the cochlea. The reduction in these macrophages in RC was not statistically significant (p=0.5981; 30.5 vs 15.674 cells/mm^3^). At 336 days post-implantation, mDex-CIs suppress the infiltration of CX3CR1+ macrophages in the ST (p<0.0001; 18.725 vs 0.872 cells/mm^3^), RC (p=0.0202; 38.05 vs 7.782 cells/mm^3^), and LW (p<0.0001; 45.425 vs 9.834 cells/mm^3^) in the basal turn of the cochlea. A subset of these infiltrating CX3CR1+ macrophages is MHCII+CX3CR1+ antigen-presenting macrophages.

**Fig. 3.**
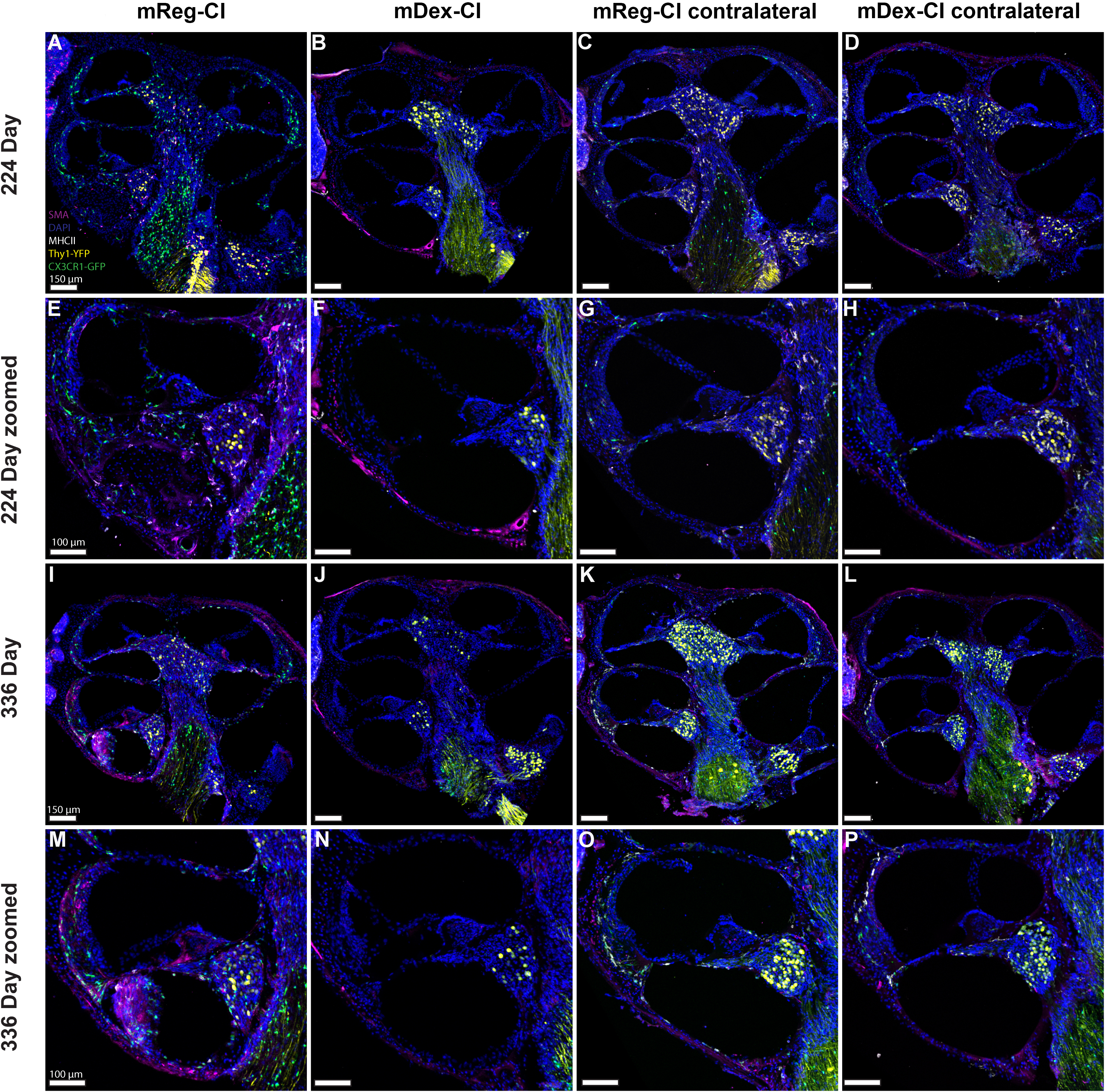
Dexamethasone-eluting implant reduces the density of CX3CR1+ macrophages, density of MHCII+CX3CR1+ macrophages, and volume of α-SMA in the basal turn of the cochlea. CX3CR1+/eGFP Thy1+/eYFP mice were implanted with either mReg-CI or mDex-CI and euthanized at 224 days (A-H) or 336 days (I-P) post-implantation. Maximum intensity z-projections of 3D confocal image stacks taken from 30-μm-thick mid-modiolar sections showing the basal turn of the cochlea from implanted and non-implanted, contralateral ears.

**Fig. 4.**
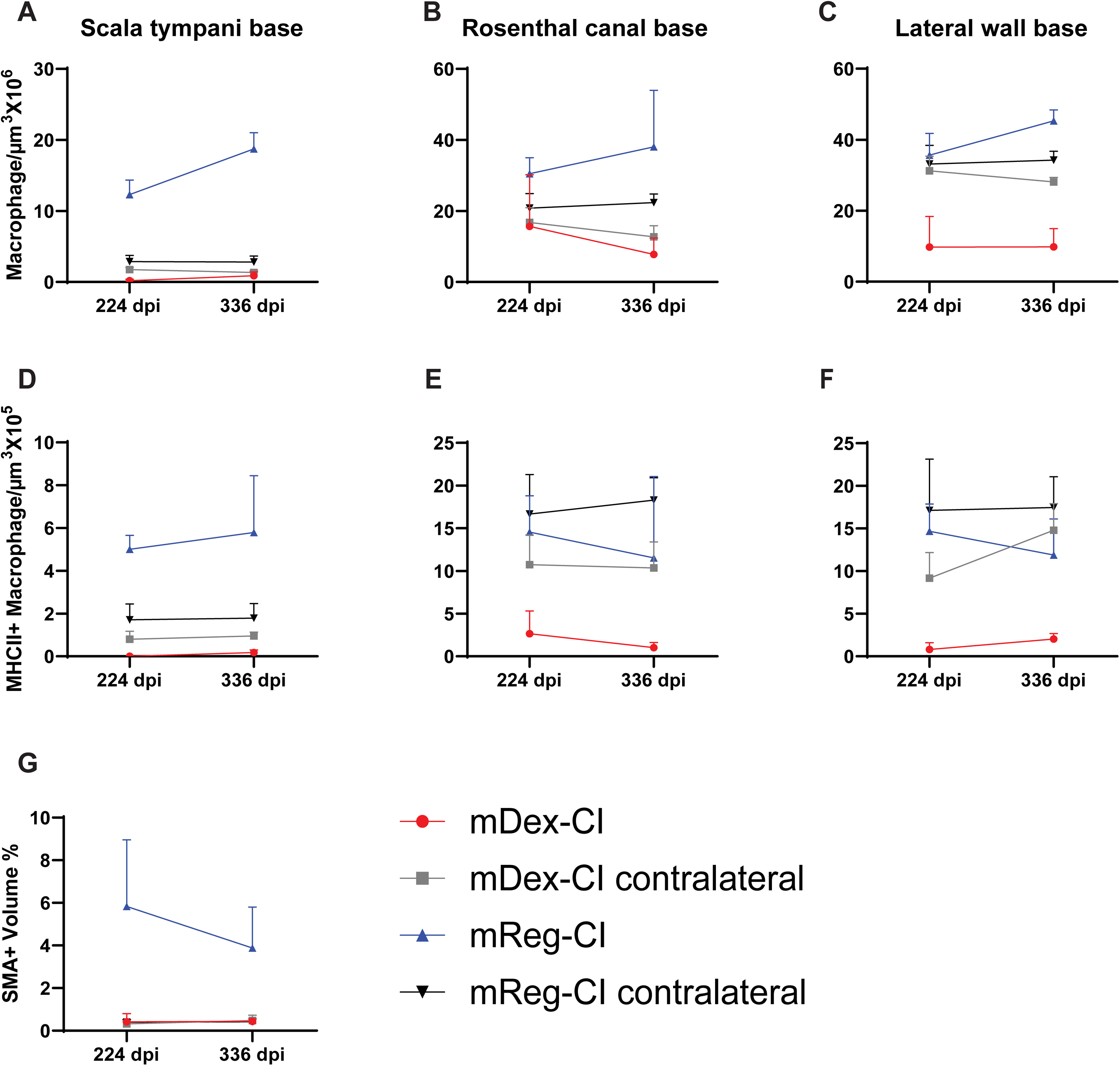
Quantification of CX3CR1+ macrophage density **(A-C)**, MHCII+CX3CR1+ macrophage density **(D-F)**, and percentage of α-SMA+ volume occupying the scala tympani **(G)** in the basal cochlear turn following cochlear implantation. Cochlear implantation with an mReg-CI leads to an infiltration of CX3CR1+ macrophages and MHCII+CX3CR1+ macrophages, as well as fibrosis when compared with unimplanted contralateral cochleae in all measured regions at both time points. mDex-CI reduces CX3CR1+ and MHCII+CX3CR1+ macrophage infiltration when compared to mReg-CI in all measured regions at both time points. For each cochlea, the average density was calculated from three mid-modiolar sections. Error bars represent SEM.

Compared to mReg-CIs, mDex-CIs suppress the infiltration of these MHCII+CX3CR1+ cells at 224 (p=0.0041; 5.007 vs 0 cells/mm^3^) and 336 (p=0.0003; 5.791 vs 0.182 cells/mm^3^) days post-implantation in the ST; this trend was not significant in the LW (224 days: p=0.165; 14.67 vs 0.793 cells/mm3; 336 days: p=0.342; 11.89 vs 2.02 cells/mm^3^) and RC (224 days: p=0.305; 14.56 vs 2.66 cells/mm3; 336 days: p=0.317; 11.52 vs 1.02 cells/mm^3^). Following mReg-CI implantation, an α-SMA+ fibrotic response is observed in the ST. Compared to mReg-CI, mDex-CI showed significantly reduced (p=0.0070) volume of α-SMA+ fibrotic tissue in the ST at 224 days post-implantation.

This trend approached significance (p=0.0540) at 336 days post-implantation with mDex-CI showing an 88.7% reduction in α-SMA+ fibrotic tissue relative to mReg-CI (3.884% vs 0.438% volume of ST occupied), suggesting either maintained antifibrotic activity or stabilization of fibrotic tissue volume with continued drug elution. Movat’s Pentachrome staining (**Supplementary Fig. 1**) qualitatively showed robust fibrosis and neo-ossification within the scala tympani in all mReg-CI cases at 224 and 336 days, which was reduced or absent in all non-translocated mDex-CI recipients.

Unexpectedly, we observed a trend toward reduced macrophage infiltration in the contralateral (unimplanted) cochleae of mDex-CI animals compared to mReg-CI contralateral ears (**Figs. 3, 4**) in all regions, being most pronounced in the RC in the base of the cochlea (224 days: 20.78 vs 16.76 cells/mm^3^; 336 days: 22.37 vs 12.75 cells/mm^3^), though these differences did not reach statistical significance (p=0.366-0.968). These data suggest possible systemic or cross-cochlear immunomodulatory effects of locally eluted dexamethasone.

### 3.3 Dexamethasone-eluting CIs and CIs show similar neural survival

Thy1+ SGN cell bodies within RC were counted on serial mid-modiolar sections. The basal turn was separately quantified from the middle and apical turn, which are reported together as the “apex”. Counts from each individual cochlea were normalized to the contralateral ear, reporting a percentage of neurons remaining relative to the contralateral counts. As shown in **Fig. 5**, when normalized to their contralateral ear, cochleae implanted with mDex-CIs and mReg-CIs did not show a significant difference in neuron density across the entire cochlea at both time points (p=0.529-0.908).

**Fig. 5.**
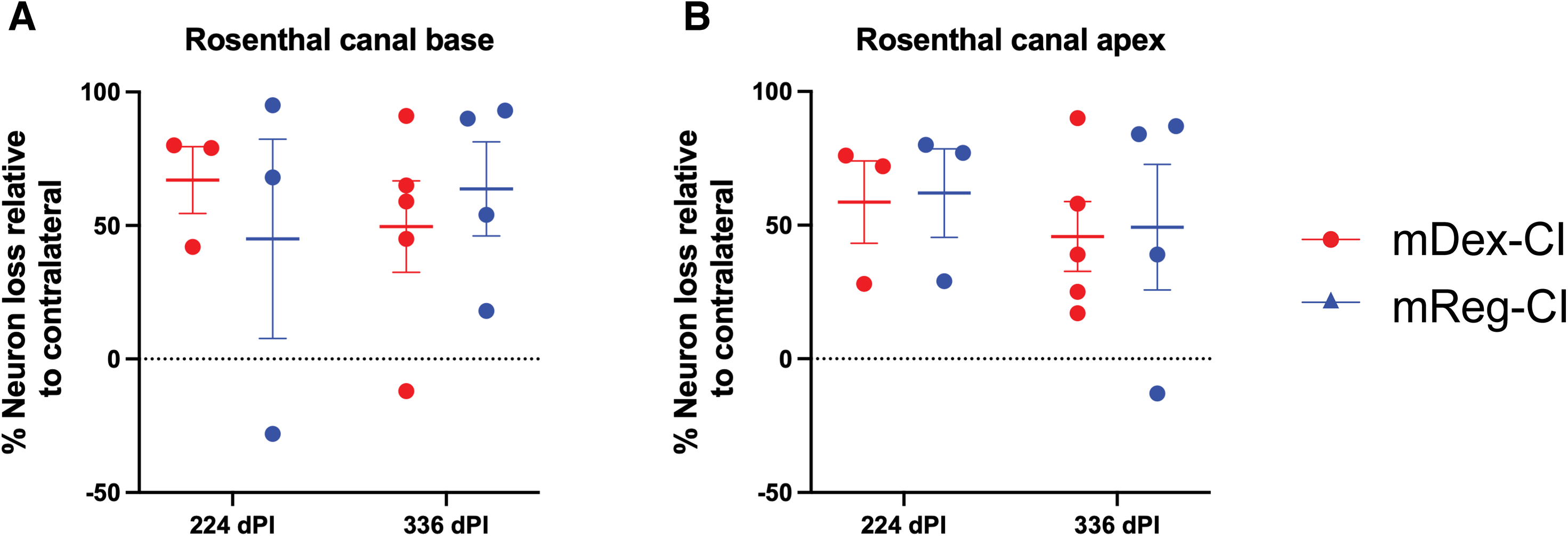
Quantification of Thy1+/eYFP neuron survival in RC in the basal turn (**A)** and apical turn **(B)** of the cochlea. There is not a significant difference between neuron survival when comparing mDex-CI to mReg-CI at either 224 or 336 days post-implantation at the apical or basal RC. Neurons were counted in implanted cochlea, and counts were referenced against neuron counts in unimplanted contralateral cochlea of the same subject to reach a measurement of percent survival. Cells were counted using IMARIS image analysis software from images taken from three consecutive mid-modiolar sections for each sample. Error bars represent SEM.

### 3.4 Translocation of CIs

Translocation through the basilar membrane was observed in 8 total cases (tabulated in **Table 1** and shown in **Fig. 6**). When compared to a non-translocated mReg-CI, translocated mReg-CIs result in significantly increased CX3CR1+ macrophage infiltration in the ST (p=0.0359). In the presence of translocation, mDex-CIs suppress the infiltration of CX3CR1+ macrophages in the ST compared to a translocated mReg-CI at the same time point (p=0.0039) (**Fig. 7**). The reduction in MHCII+CX3CR1+ macrophages was not statistically significant. In mDex-CI cochleae with translocation, histological analysis revealed robust fibrotic and osseous responses strictly localized to the site of basilar membrane breach (scala media and scala vestibuli adjacent to the electrode), while the remaining scala tympani was free of fibrosis and neo-ossification (**Fig. 6E-F**). In contrast, mReg-CI translocations produced widespread fibrosis and neo-ossification extending throughout the basal ST, including regions distant from the translocation site (**Fig. 6A-D**). This pattern suggests that mDex-CI permits localized wound-healing responses while preventing generalized foreign body reactions. No data are available for translocated mDex-CIs at 336 days post-implantation because no translocation occurred in this group.

**Fig. 6.**
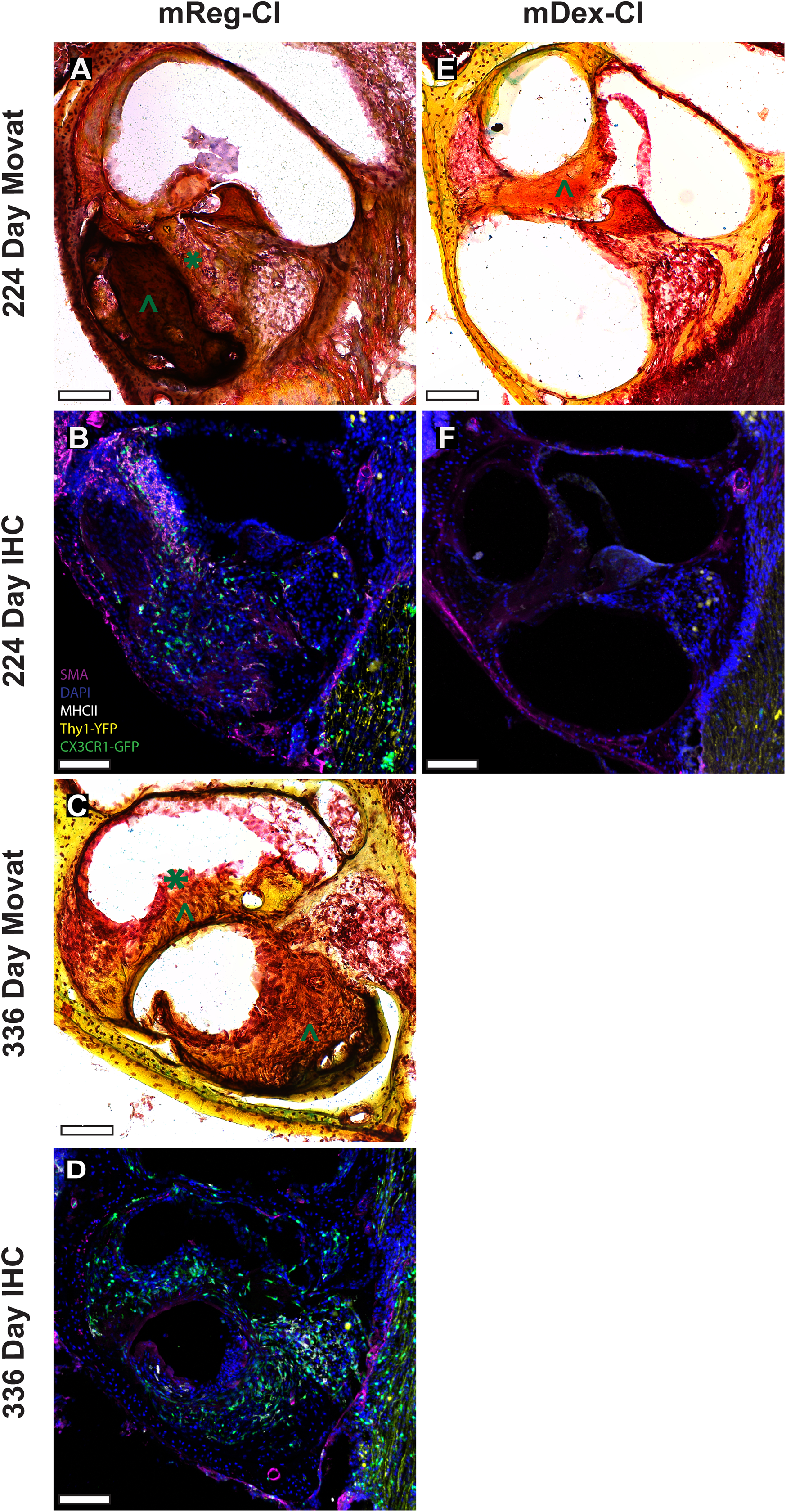
**Typical mid-modiolar sections** are shown from translocated mReg-CI (**A, B, C, D**) and mDex-CI (**E, F**). In the event of an electrode translocating through the basilar membrane, mDex-CI reduces the density of CX3CR1+ macrophages and MHCII+CX3CR1+ macrophages in the basal turn of the cochlea when compared with an mReg-CI with translocation. Despite the reduction in macrophages, there remains a strong reactive fibrotic response and neo-ossification localized to the site of translocation in mDex-CI, with an otherwise open scala tympani, as shown by Movat’s Pentachrome staining in (**E**). In the presence of a translocation, mReg-CI cochleae showed robust fibrosis and neo-ossification spanning multiple scalae (**A, C**). Areas of fibrosis and neo-ossification are marked with green “*” and “^”, respectively. Scale bars are 100 μm.

**Fig. 7.**
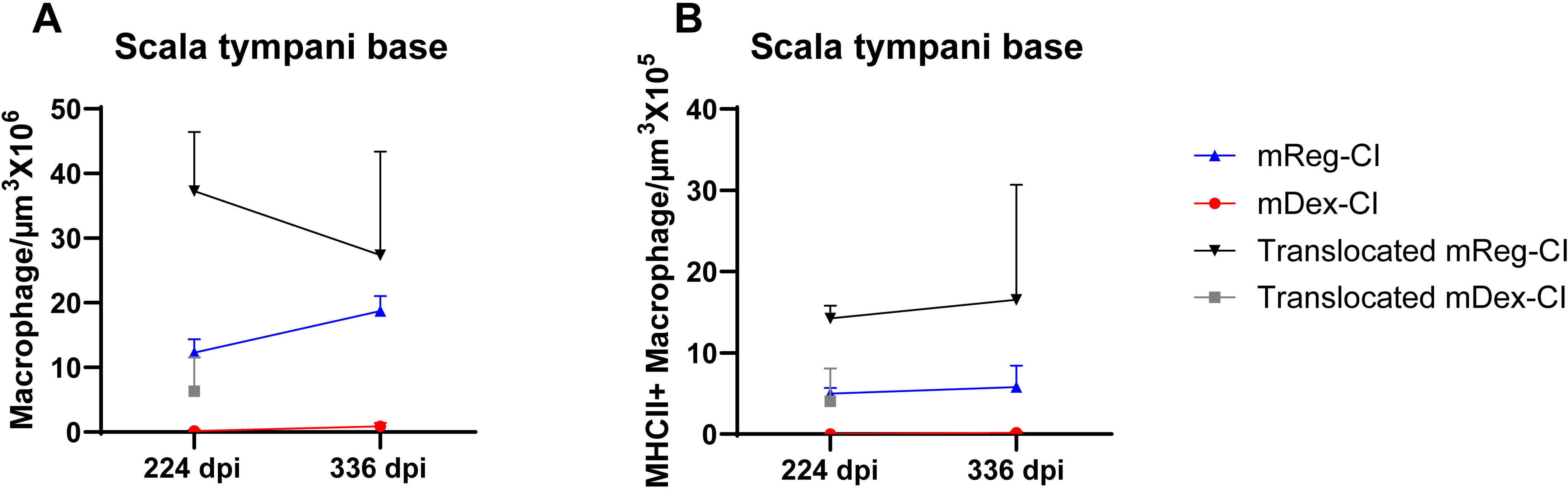
Macrophage infiltration in translocated vs non-translocated cases. Quantification of CX3CR1+ macrophage density **(A)** and MHCII+CX3CR1+ macrophage density **(B)** in the basal cochlear turn following cochlear implantation in translocated and non-translocated CI cases. In the event of translocation, mDex-CI reduces CX3CR1+ macrophage density and MHCII+CX3CR1+ macrophage density in the basal turn of the cochlea. For each cochlea, the average density was calculated from three mid-modiolar sections.

**Table 1.**
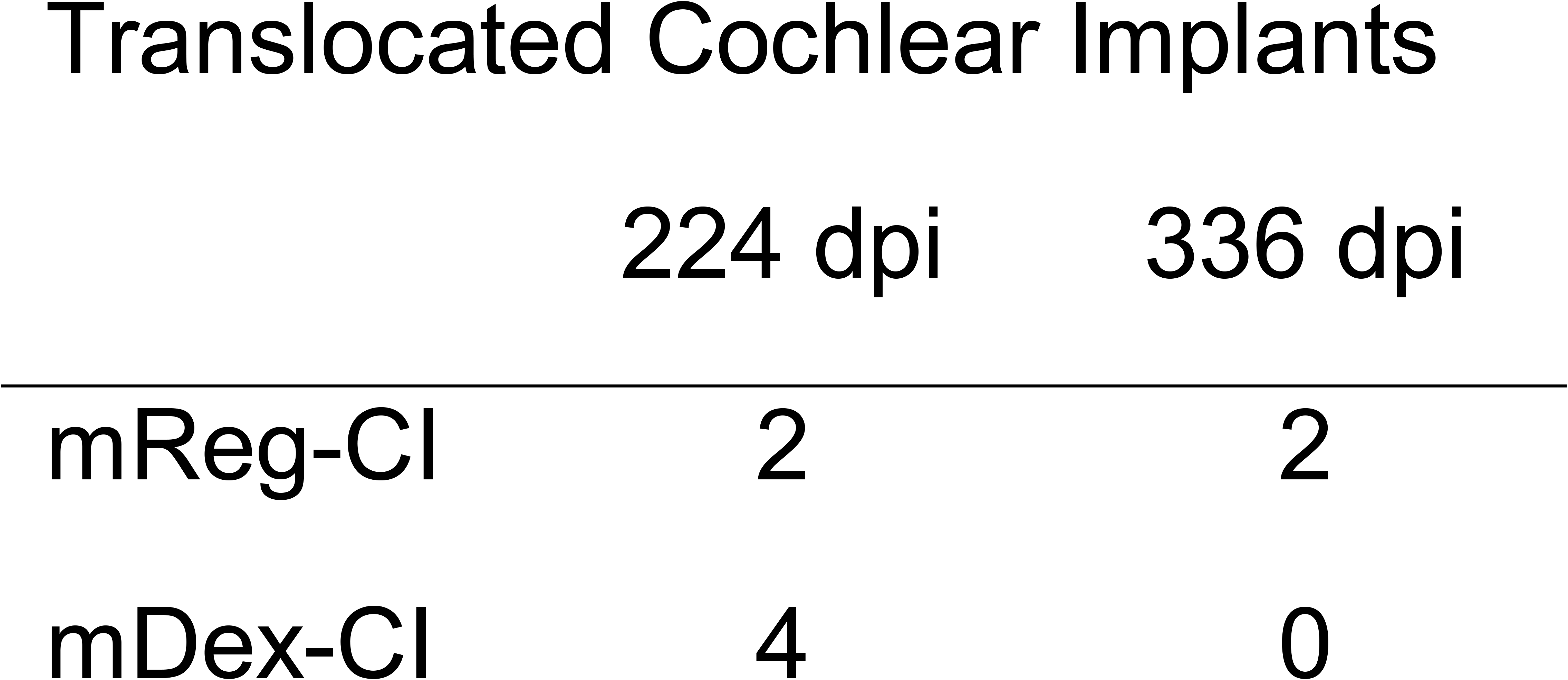
The number of cochleae (n) in each group and time point where the electrode array had translocated through the basilar membrane of the basal turn rather than resting in the ST.

## 4. Discussion

The data demonstrate persistent macrophage infiltration, fibrotic tissue deposition, and neo-ossification in the mouse cochlea after mReg-CI out to 336 days post-implantation. mDex-CI mitigated this immune cell infiltration and tissue response for at least 336 days post-implantation, in the setting of continuous low-level dexamethasone elution. These data extend the findings of Rahman et al. (2025) showing mDex-CI suppression of the FBR from 112 days out to 336 days post-implantation. This long-term suppression of the FBR with mDex-CI is consistent with published human data (Briggs et al., 2020) showing that dexamethasone-eluting cochlear implants are associated with lower electrode impedances out to 24 months post-CI.

Low-level dexamethasone elution was sufficient to suppress cochlear macrophage populations to levels approaching unimplanted contralateral ears through 336 days post-implantation, demonstrating that sustained local glucocorticoid delivery can durably reshape the intracochlear immune environment well into the chronic phase of the FBR. Although macrophage infiltration and intracochlear fibrosis are mechanistically linked following cochlear implantation, the antifibrotic effect of mDex-CI likely reflects broader anti-inflammatory activity rather than macrophage suppression and reduced antigen presentation in isolation. Glucocorticoids are known to attenuate lymphocyte recruitment, suppress pro-inflammatory cytokine production, and directly inhibit fibroblast proliferation and collagen deposition, any of which may contribute to the reduced fibrotic response observed here (Cain and Cidlowski, 2017; Weiner et al., 1987; Slavin et al., 1994). Consistent with a multi-pathway mechanism, we have previously shown that isolated macrophage depletion in the setting of cochlear implantation is insufficient to suppress the post-CI tissue response (Rahman et al., 2023). Together, these observations indicate that the mitigative effects of mDex-CI on the FBR extend beyond suppression of macrophage infiltration, with macrophage modulation representing one component of a broader immunomodulatory effect.

An unexpected finding of this study was the trend toward reduction in CX3CR1+ macrophages and MHCII+CX3CR1+ antigen-presenting cells in the contralateral, unimplanted cochleae of animals receiving an mDex-CI relative to the contralateral cochleae of mReg-CI animals. Because the implant was placed unilaterally, suppression of the immune response in the opposite ear suggests that locally eluted dexamethasone exerts an effect beyond the implanted cochlea. Several mechanisms could account for this observation. First, dexamethasone eluted into the scala tympani may be absorbed into the systemic circulation and redistributed to the contralateral inner ear, where even low concentrations of glucocorticoid may be sufficient to dampen monocyte recruitment and antigen-presenting macrophage activation. Second, the two cochleae communicate with a shared cerebrospinal fluid compartment through the cochlear aqueduct, providing an anatomic route by which intracochlear dexamethasone could reach the perilymph of the contralateral ear. Third, cochlear implantation provokes a systemic inflammatory response, and dexamethasone may blunt this circulating inflammatory signal, thereby reducing the priming and recruitment of immune cells to the contralateral cochlea. Importantly, this contralateral immunomodulatory effect of mDex-CI may prove beneficial in the setting of sequential hearing preservation surgery, where the second ear may show a more rapid rise in electrode impedances (Flom et al., 2025) and greater risk of loss of residual acoustic hearing (Kocharyan et al., 2025). Further studies measuring systemic and contralateral perilymph dexamethasone concentrations are needed to define the route and clinical significance of this cross-cochlear effect.

Although there was a strong inhibition of the CI FBR with mDex-CI, it remained permissive of local reparative responses, including fibrosis and ossification seen directly adjacent to sites of basilar membrane translocation. Notably, macrophage infiltration remained suppressed at sites of local repair of translocation in mDex-CI, suggesting this process may not be dependent on the innate immune response. This would reflect a clinically favorable profile for a dexamethasone-eluting CI in which it reduces the cochlear FBR but does not inhibit beneficial reparative responses, such as repair of scalar translocation or round window sealing. This finding also suggests a complementary role for other surgical adjuncts, including robotic CI insertion (Claussen et al., 2022b; Khan et al., 2025; Khan et al., 2026b) in mitigating other factors (e.g., surgical trauma) that may negatively impact CI outcomes.

We observed that CI was associated with loss of neural cell populations, similar to the neural degeneration seen in prior studies of mouse CI (Wu et al., 2026). The level of neural degeneration was not significantly different between mDex-CI and mReg-CI. However, the study design was limited in assessment of neural survival in several aspects. Thy1+ may not reliably mark all SGN cell bodies (Feng et al., 2000; Taylor-Clark et al., 2015). Additionally, counting of select mid-modiolar sections may produce counts that are biased by histologic sectioning orientation and exclude edge portions of the sectioning series. Finally, our density counts were limited to SGN cell bodies and did not assess the peripheral or central processes, synaptic connections, or myelination status of the auditory nerve.

A limitation of this study is that no electric stimulation was delivered. We have previously published that electric stimulation did not have an effect on the innate immune response within the first 3 weeks in a mouse CI model (Claussen et al., 2022a). Further, the current generation of mouse CIs is prone to short-circuiting beyond one month of implantation, prohibiting an experimental design that would allow reliable electric stimulation for 224-336 days post-implantation. An additional limitation was our use of normal-hearing mice, which do not represent the typical CI population. Further work is necessary to assess whether auditory injury of varied etiology (e.g., noise versus ototoxic) alters the CI FBR by priming the local cochlear immune response (Hirose et al., 2005; Wood and Zuo, 2017; Kaur et al., 2018).

In summary, the data presented here demonstrate a long-term reduction in the local immune and “tissue” response after dexamethasone-eluting cochlear implantation. This effect was associated with continued low-level drug elution; thus, we cannot conclude if this outcome persists after exhaustion of dexamethasone elution. Future studies are needed in both pre-clinical models and long-term human dexamethasone-eluting CI follow-up to assess if these effects persist after total completion of dexamethasone elution.

## Data availability

The datasets generated and analyzed in this study will be made publicly available upon acceptance.

## Ethics statement

All animal procedures were approved by the Institutional Animal Care and Use Committee (IACUC) of the University of Iowa (Protocol No. 4021302) and were performed in accordance with institutional guidelines.

## CRediT authorship contribution statement

Aditya Alluri: Investigation, Formal analysis, Visualization, Writing – original draft, Writing – review & editing. Bryce Hunger: Investigation, Formal analysis, Visualization, Writing – original draft, Writing – review & editing. Fahad Hossain: Investigation, Writing – review & editing. Shakila Mahmuda Fatima: Investigation, Writing – review & editing. Muhammad Taifur Rahman: Investigation, Writing – review & editing. Brian J. Mostaert: Conceptualization, Investigation, Writing – review & editing. Robert Gay: Conceptualization, Resources, Writing – review & editing. Ya Lane Enke: Conceptualization, Resources, Writing – review & editing. Marlan R. Hansen: Conceptualization, Formal analysis, Supervision, Funding acquisition, Writing – review & editing. Alexander D. Claussen: Conceptualization, Formal analysis, Supervision, Funding acquisition, Writing – review & editing.

## Acknowledgments

We thank Lynn Teesch for performance of LC–MS/MS studies.

---

**Supplementary Fig. 1.** Typical mid-modiolar sections stained with Movat’s Pentachrome to highlight neo-ossification and fibrosis are shown from non-translocated mReg-CI (**A, E**), non-translocated mDex-CI (B, F), and contralateral ears (C). Robust fibrosis and neo-ossification are observed in mReg-CI, mostly confined to the scala tympani. mDex-CI showed no fibrosis or neo-ossification in the scala tympani at 224 days (**B**) and only sparse fibrosis in the basal scala tympani at 336 days (**F**). Areas of fibrosis and neo-ossification are marked with red “*” and “^”, respectively. Scale bars are 100 μm.

