## Supplementary figures and images for "Long-term mitigation of the foreign-body response with dexamethasone-eluting cochlear implants in mice"

### Supplemental Table 1

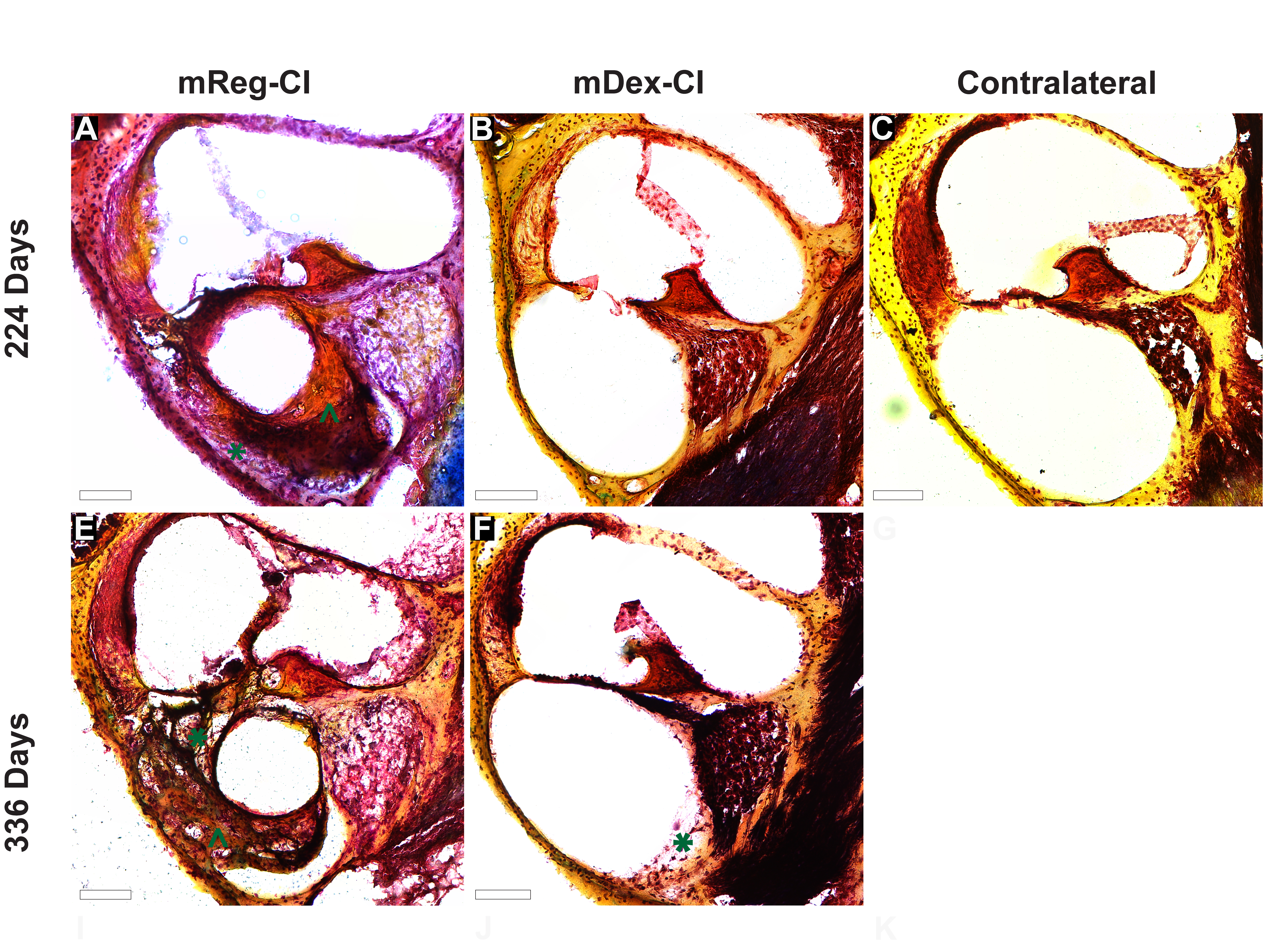
